# Transient dynamics in plant-pollinator networks: Fewer but higher quality of pollinator visits determines plant invasion success

**DOI:** 10.1101/2022.05.03.490461

**Authors:** Fernanda Valdovinos, Sabine Dritz, Robert Marsland

## Abstract

Invasive plants often use mutualisms to establish in their new habitats and tend to be visited by resident pollinators similarly or more frequently than native plants. The quality and resulting reproductive success of those visits, however, have rarely been studied in a network context. Here, we use a dynamic model to evaluate the invasion success and impacts on natives of various types of non-native plant species introduced into thousands of plant-pollinator networks of varying structure. We found that network structure properties did not predict invasion success, but non-native traits and interactions did. Specifically, non-native plants producing high amounts of floral rewards but visited by few pollinators at the moment of their introduction were the only plant species able to invade the networks. This result is determined by the transient dynamics occurring right after the plant introduction. Successful invasions increased the abundance of pollinators that visited the invader, but the reallocation of the pollinators’ foraging effort from native plants to the invader reduced the quantity and quality of visits received by native plants and made the networks slightly more modular and nested. The positive and negative effects of the invader on pollinator and plant abundance, respectively, were buffered by plant richness. Our results call for evaluating the impact of invasive plants not only on visitation rates and network structure, but also on processes beyond pollination including seed production and recruitment of native plants.

## Introduction

Species invasions are one of the six global change drivers threatening biodiversity worldwide (Tylianakis et al. 2008). Plants make up the largest and most studied group of invasive species globally (Pyšek et al. 2008, Downey and Richardson 2016), which often use mutualisms to establish in their new habitats (Richardson et al. 2000, Traveset and Richardson 2014, Parra-Tabla and Arceo-Gómez 2021). In particular, the interaction of non-native plants with resident pollinators (native or non-native) plays an important role in the reproductive success of invasive plants (Ghazoul 2002, Traveset and Richardson 2014, Parra-Tabla and Arceo-Gómez 2021). Studies analyzing the interactions of non-native plants within plant-pollinator networks indicate that these species are well-integrated into the networks by showing that they share flower visitors with native plants (Aizen et al. 2008, Bartomeus et al. 2008, Kaiser-Bunbury et al. 2011, Traveset et al. 2013, Montero-Castaño and Vilà 2017) or that they are visited either similarly or more frequently than the natives (Lopezaraiza–Mikel et al. 2007, Montero-Castaño and Vilà 2017, Parra-Tabla et al. 2019, Seitz et al. 2020). However, the long-term persistence of pollinator-dependent plants in their new community not only depends on receiving pollinator visits but also on the pollinators’ efficiency in transporting their conspecific pollen and the subsequent plant reproduction (Parra-Tabla and Arceo-Gómez 2021).

The effects of these two key factors (i.e., pollinator efficiency and plant reproductive success) on pollinator-dependent plant invasions have been rarely studied in the context of plant-pollinator networks (Parra-Tabla and Arceo-Gómez 2021). Some findings suggest that a non-native plant receiving many pollinator visits will not necessarily persist in its new community because those visits might not contribute to its reproduction success. De Santiago-Hernandez et al. (2019) found that only 59% of floral visitors contribute to seed production. Indeed, non-native plants receiving few but high quality visits can also persist in their new community. Thompson and Knight (2018) show that non-native plants can exhibit high reproductive success when visited by only one or a few pollinator species. In contrast, other studies find that several invasive species exhibit generalized floral traits (Parra-Tabla and Arceo-Gómez 2021), are visited by many and abundant pollinator species (Bartomeus et al. 2008, Vilà et al. 2009), and tend to be network hubs (Albrecht et al., 2014). These contrasting empirical patterns have been obtained for plant species that had already invaded the networks and do not necessarily explain their invasion success from the early stages of their introduction.

Our understanding of the critical, early stages that determine the success of a species invasion can greatly benefit from studying the transient dynamics right after a new species is introduced into a community. The increasing recognition that many ecological phenomena occur before the system reaches an equilibrium has called for theory focusing on transient as opposed to equilibrium dynamics (Hastings et al. 2018, 2021, Morozov et al. 2020, Francis et al. 2021, Abbott et al. 2021). Dynamical transients are defined as the non-asymptotic dynamical regimes that persist for less than one to ‘as many as tens of generations’ (Hastings et al. 2018). Computer simulations of network dynamic models can help us understand the transient dynamics that occurs within a community after a species introduction, and be used to evaluate whether non-native traits and network characteristics predict the invasion success of the introduced species.

Invasive plants can affect plant-pollinator communities negatively by competing with native plants for pollinators or by increasing heterospecific pollen transfer (Traveset and Richardson 2006, 2014, Morales and Traveset 2009, Arceo-Gómez and Ashman 2016, Kaiser-Bunbury et al. 2017, Parra-Tabla et al. 2021), but also have null (Kaiser-Bunbury et al. 2011) or even positive effects on the communities via increased abundance of native pollinators (Lopezaraiza–Mikel et al. 2007, Bartomeus et al. 2008, Carvalheiro et al. 2008, Valdovinos et al. 2009). These plants can also affect the networks’ structure by modifying the strength (Kaiser-Bunbury et al. 2017) and number (Bartomeus et al. 2008, Valdovinos et al. 2009) of species interactions, the natives’ position within the network (Aizen et al. 2008, Albrecht et al. 2014), and network-level metrics such as modularity, nestedness, or connectance (Bartomeus et al. 2008, Valdovinos et al. 2009). However, the mechanisms behind those network changes and the impacts of those network changes on the native species are not entirely understood (Parra-Tabla and Arceo-Gómez 2021).

Here, we use a dynamic plant-pollinator network model to evaluate the efficiency of pollinator visits non-native plants receive and their resulting reproductive success at the critical early stages of invasion. In addition, we determine their impact on native species’ reproductive success at equilibrium. In terms of non-native traits, we focus on rewards production, pollen attachability, and level of generality (i.e., number of pollinator species visiting them) because these are highly variable traits that influence the reproductive success of pollinator-dependent plants (Olesen et al. 2011, Baude et al. 2016, Timberlake et al. 2019, Filipiak et al. 2022). We answer three questions: 1) How does higher reward production, pollen attachability, and number of pollinator visitors affect the reproductive success of non-native plants? 2) How does the quantity and quality of visits a plant receives from resident pollinators affect their invasion success? 3) How do plant invasions impact network structure and the reproduction success of native plants?

## Materials and methods

### Binary vs. weighted network structures

The binary structure of networks represents species as nodes and their interactions as binary links, while the weighted structure provides information about the strength of those interactions as weighted links. We use the visitation rate of each pollinator species to each plant species (function *V*_*ij*_ in Table 1) to determine the weighted structure, which depends on the abundances of plant and pollinator species, the pollinators’ foraging efforts, and visitation efficiency. Empirical studies most often use this definition of weighted structures because frequency of visits is what researchers most often record in the field (e.g., Bartomeus et al. 2008, Vilà et al. 2009, Kaiser-Bunbury et al. 2011, 2017). We used the 1200 binary structures from Valdovinos et al. (2018), composed of three sets of 400 networks centered at three combinations of richness (S) and connectence (C), with values: S = 40 and C = 0.25, S = 90 and C = 0.15, and S = 200 and C = 0.06. These combinations represent three points in the empirically observed relation between richness and connectance, and recreate structural patterns of empirically observed networks including their heterogenous degree distribution and nestedness. Half of the networks at each set are nested and the other half, non-nested, with NODFst values ranging between -0.33 and 2.3. These networks maintain the empirically observed mean ratio of animal to plant species of 2.5 (Jordano et al. 2003). The weighted structures emerged from the network dynamics (see below).

**Table 1.** Biological processes and assumptions in Valdovinos et al.’s (2013) model.

| Biological process | In the model | Assumption |
| --- | --- | --- |
| Visitation rate | $V_{ij} = \alpha_{ij}\tau_j a_j p_i$ | Depends on the pollinator $j$'s foraging effort ($\alpha_{ij}$) assigned to plant $i$, $j$'s flying efficiency ($\tau_j$), and the plant ($p_i$) and pollinator ($a_j$) densities. |
| Pollination events | $\sigma_{ij}V_{ij}$ | Only a fraction of pollinator visits of $j$ to $i$ ($V_{ij}$) produce pollination events, determined by the proportion of conspecific pollen carried by the pollinator (visit quality function $\sigma_{ij}$). |
| Total pollination events | $\sum_{j \in A_i} \sigma_{ij}V_{ij}$ | Pollination events summed over all the pollinator species visiting the plant (set $A_i$). |
| Seed production | $e_i \sum_{j \in A_i} \sigma_{ij}V_{ij}$ | Only a fraction of the total pollination events become seeds, determined by the seed production efficiency of the plant species (parameter $e_i$). |
| Seed recruitment | $\gamma_i e_i \sum_{j \in A_i} \sigma_{ij}V_{ij}$ | Only a fraction of seeds produced recruit to adults, determined by the competition among plants (function $\gamma_i$). |
| Consumption of rewards | $V_{ij} b_{ij} \frac{R_i}{p_i}$ | In each visit, pollinators consume a fraction of the floral rewards offered by the plant individual ($R_i/p_i$) at a rate $b_{ij}$. |
| Recruitment to adult pollinators | $c_j \sum_{i \in P_j} V_{ij} b_{ij} \frac{R_i}{p_i}$ | Floral rewards consumed by the pollinator species summed over all the plant species the pollinator species visits (set $P_j$) are converted into new pollinator adults at a rate $c_j$. |
| Production of rewards | $\beta_i p_i - \varphi_i R_i$ | Floral rewards of a plant population increase with its population density in a saturating manner, with rewards production decelerating as rewards increase up to the maximum of $\beta_i p_i / \varphi_i$ when the rewards production stops. |
| Adaptive foraging | Equation (4) | A pollinator increases its foraging effort to plants with more rewards, by reassigning its efforts from plants with fewer rewards. Foraging efforts are fractions that can take a maximum value of 1 (the pollinator assigns all its effort to that plant) and they sum to 1 over all plants the pollinator visits. |
| Efforts of a fixed forager | $1/k_{aj}$ | Pollinators without adaptive foraging are assumed to have fixed foraging efforts across all the plants they visit equal to one over the number of plant species the pollinator visits |

### Network dynamics

We used Valdovinos et al.’s (2013) model, which assumes that all plant species in the network depend on animal pollination for reproduction to simulate the network dynamics.

Several previous studies have used and analyzed this model (Valdovinos et al. 2013, 2016, 2018, Valdovinos and Marsland 2020), including its sensitivity to parameter values. We summarize the biological processes encapsulated in the model and its assumptions in Table 1, provide the definitions and values of its functions and parameters in Table 2, and analyze the robustness of our results across parameter values in Appendix S1 (Online Supplementary Information). This model defines the population dynamics (over time *t*) of each plant (Eq. 1) and pollinator (Eq. 2) species of the network, as well as the dynamics of floral rewards (Eq. 3) of each plant species, and the foraging effort (Eq. 4) that each pollinator species (per-capita) assigns to each plant species as follows:

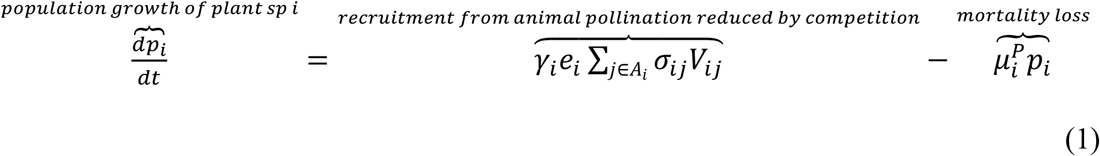

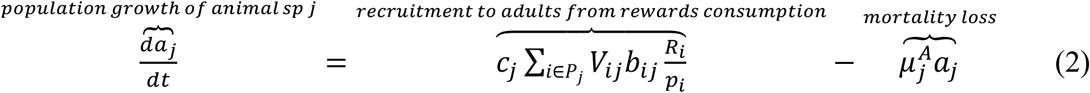

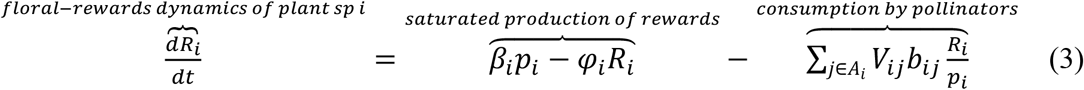

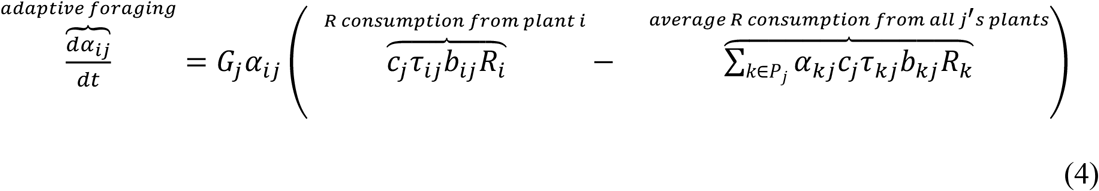

Previous work used this model to evaluate the invasion success and impacts of non-native pollinators on plant-pollinator networks (Valdovinos et al. 2018). However, the dynamics of pollinators and plant in this model are very different. That is, the equations describing their population dynamics encapsulate biological mechanisms that differ drastically between pollinators and plants (see Eqs. 1 and 2; Table 1), which results in very different dynamical outputs and effects on other species in the network (Valdovinos et al. 2013, 2016, 2018, Valdovinos and Marsland 2020). Moreover, these differences in modeled population dynamics may provide insights into the mechanisms influencing the invasion processes of pollinators vs. plants in ecological networks.

**Table 2.**
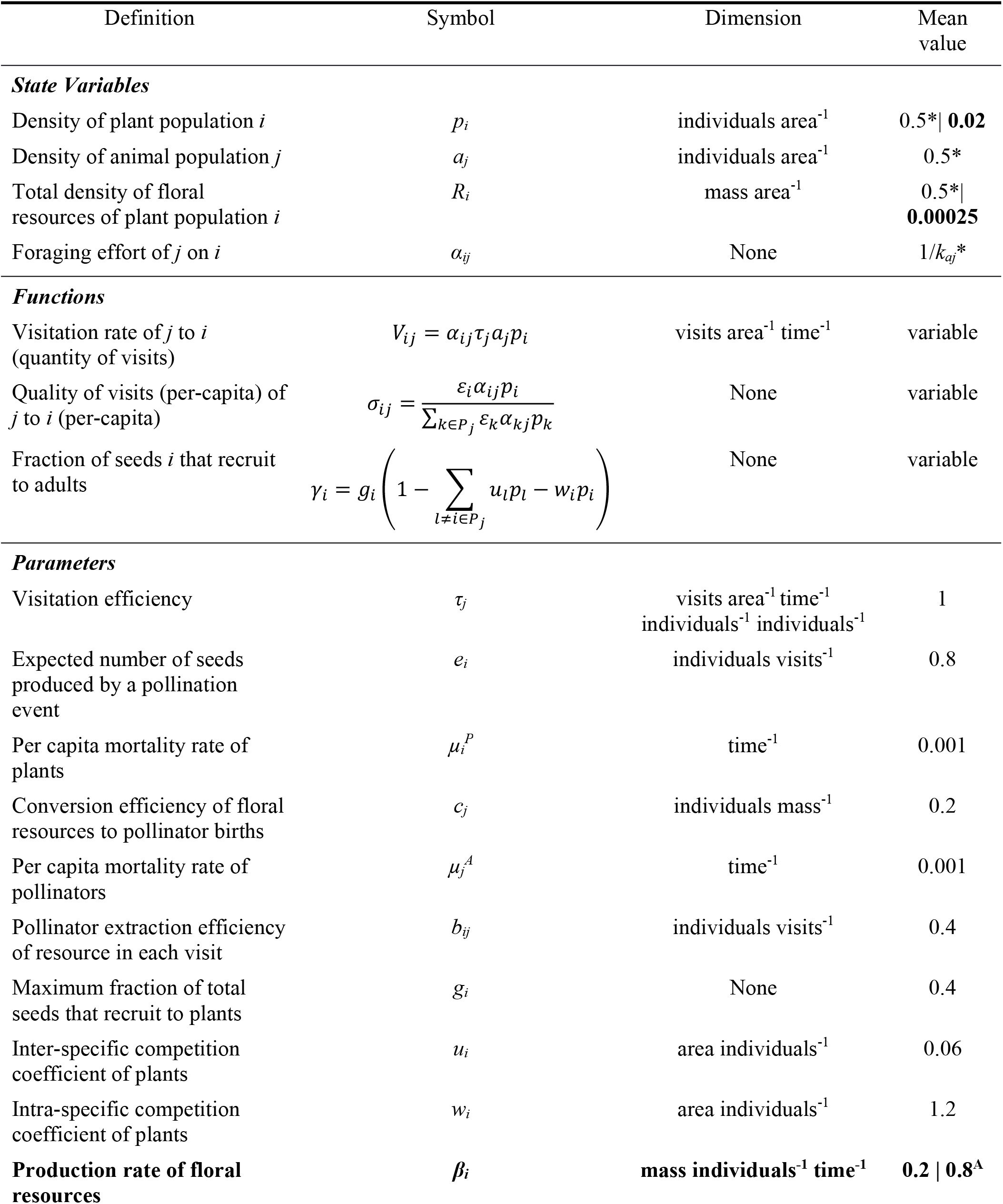
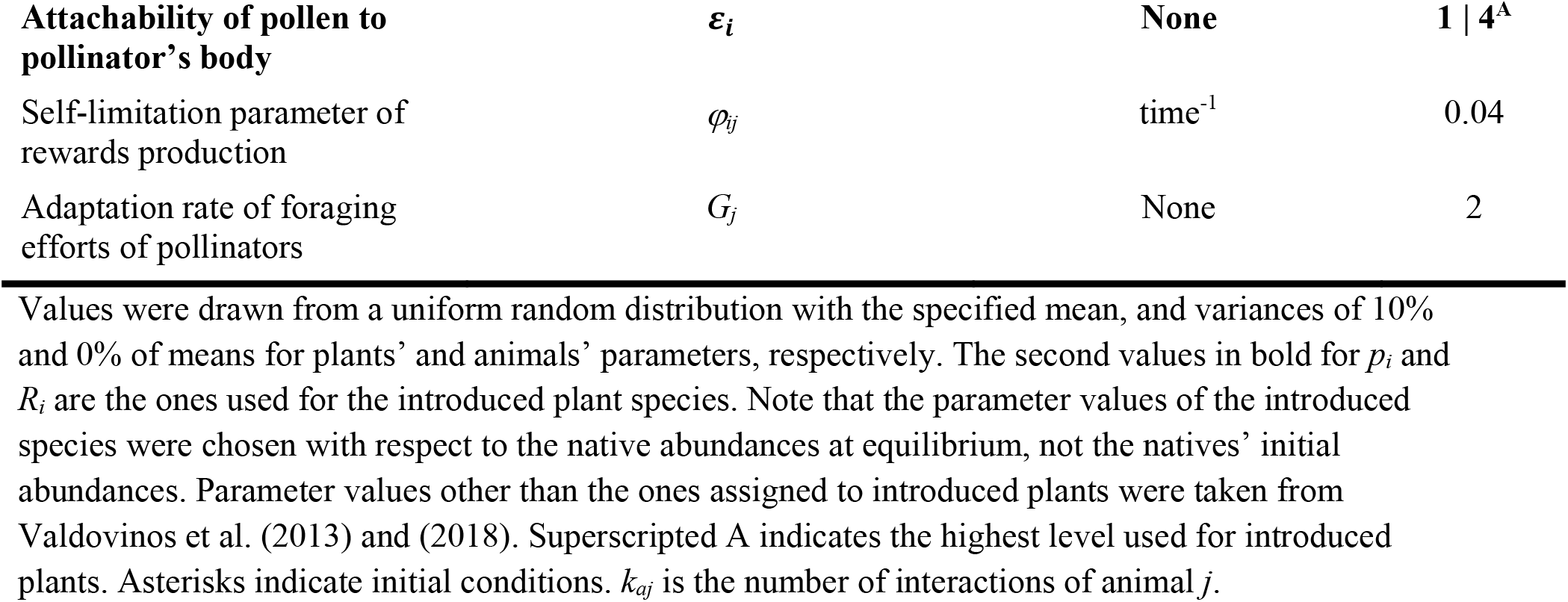
Model state variables, functions, and parameters.

We ran the model for 10,000 timesteps prior to the plant introductions and another 10,000 timesteps after the introduction. We analyzed both the transient dynamics immediately after the plant introduction (during the first 2,000 timesteps after the introduction) and the equilibrated dynamics (at 10,000 timesteps after the introduction). The simulations generally equilibrated at around 3,000 timesteps, so running them longer ensured we captured the dynamics at equilibrium.

### Non-native introductions

We introduced 8 types of plant species to each network (one per simulation) based on all combinations of two levels of three properties (see Table 3) at *t* = 10,000, with density equal to the plant extinction threshold, 0.02, and reward density 0.02 times that of the average native at equilibrium (i.e., 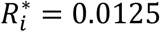, see Eq. S3) to keep the initial rewards density per plant similar between non-native and native plants. Therefore, the introduced plant species always starts out at a double disadvantage with respect to the native plants because its initial abundance *(p*_*x*_ 0.02), and the foraging effort pollinators assign to it *(a*_*xj*_ 0.0001) are very small compared to the abundance at equilibrium of native plants at the moment of its introduction (average 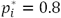) and the foraging efforts that they receive at equilibrium (average *a*_*ij*_ 0.3). The extinction threshold was set in previous work based on the Allee effect experienced by plants for the parameter values shown in Table 2 (Valdovinos et al. 2013, 2016, 2018).The pollinator species that initially visited the introduced plant were chosen randomly from: (1) all pollinator species, (2) most-generalist pollinator species, (3) most-specialist pollinator species. These three options of “linkage algorithms” are called hereafter ‘random’, ‘most connected’, and ‘least connected’, respectively. The foraging effort of native pollinators initially visiting the introduced plant was set to 0.0001 (of a total of 1 summed over all the interactions of the pollinator), which was subtracted from the highest effort of the pollinator so the effect of the effort subtraction was negligible. We conducted a total of 28,800 plant introductions (1200 networks × 8 plant types × 3 linkage algorithms).

**Table 3.** Properties of the non-native plants introduced.

| Factor (property) | Description of level 1 | Description of level 2 |
| --- | --- | --- |
| **Generality (# links)** | Specialist (*average # links of 30% most specialist natives*) | Generalist (*average # links of 30% most generalist natives*) |
| **Pollen attachability ($\epsilon_i$)** | Same as average native | Four times higher than average native* |
| **Rewards production ($\beta_i$)** | Same as average native | Four times higher than average native* |

\*We chose the high levels of pollen attachability and rewards production to be four times higher than those of the average natives, because those levels show clear effects of the properties. Different values did not change our qualitative results.

### Analysis of the simulation results

We conducted a Classification and Regression Tree (CART) analysis using the software JMP® (Version 16.0., SAS Institute Inc., Cary, NC, 1989-2021) to evaluate which network structure properties and characteristics of non-native plants contributed most to their invasion success. We used five-fold cross validation to avoid overfitting. Network structure properties included species richness (*S*), the ratio of animal to plant species, four measures of link density [connectance (*C* = *L* / A×P, where *L* is the total number of links, A the number of pollinator species, and *P* the number of plant species), links per species (L/S), links per plant species (L/P), and links per animal species (*L/A*)], four measures of degree distribution [power law exponent for plants and animals, the standard deviation of animal generality and the standard deviation of plant vulnerability defined in Williams and Martinez (2000), four measures of niche overlap (the mean and maximum Jaccard index for plants and animals], and nestedness (see Table S1).

Introduced plant properties included the generality level, pollen attachability, rewards production, and the linkage algorithm. Network structure properties and non-native traits totaled 21 contributors for the analysis.

We evaluated the effect of successful invasions (i.e., introduced plant species that persisted at high density) on natives’ persistence, density, quality and quantity of visits. These variables were measured right before the plant introduction (*t* = 10,000), during the first 2,000 timesteps after the introductions (to understand the effects on natives of the initial introduction process), and at the end of the simulation (*t* = 20,000). We evaluated the effect of plant invasions on the networks’ weighted structure by calculating the networks’ weighted nestedness and weighted modularity before and after the invasion. These metrics were calculated using the nest.smdm() and computeModules() functions, respectively, from the R package bipartite.

## Results

### How does higher reward production, pollen attachability, and number of pollinator visitors affect the reproductive success of non-native plants?

All introduced plant species either went extinct or dramatically increased their density compared to that of native plants. Thus, we characterized the result of an introduction as either invasion failure or success. We found that specialist plants with high rewards production and high pollen attachability were the most successful invaders (see “Spec High R&P” in Fig. 1), These plants invaded 95% of the times they were introduced into the networks, while the same plant type except for being generalist invaded only 15% of the times (see “Gen High R&P” in Fig. 1A). Specialist plants with high production of rewards but average pollen attachability had an invasion success of 13% (see “Spec High R” in Fig. 1A). All other plant types never invaded. Our CART analyses (Table 4, Table S1) confirm these results, showing that among the 21 factors analyzed (17 network structure properties and 4 non-native traits, see Methods), high production of rewards contributed the most to the variation in invasion success, followed by being a specialist, and finally by having high pollen attachability. Interestingly, our CART analyses ranked the contribution of network structure to invasion success very low, with less than 5% of predictive power (Table S1).

**Figure 1.**
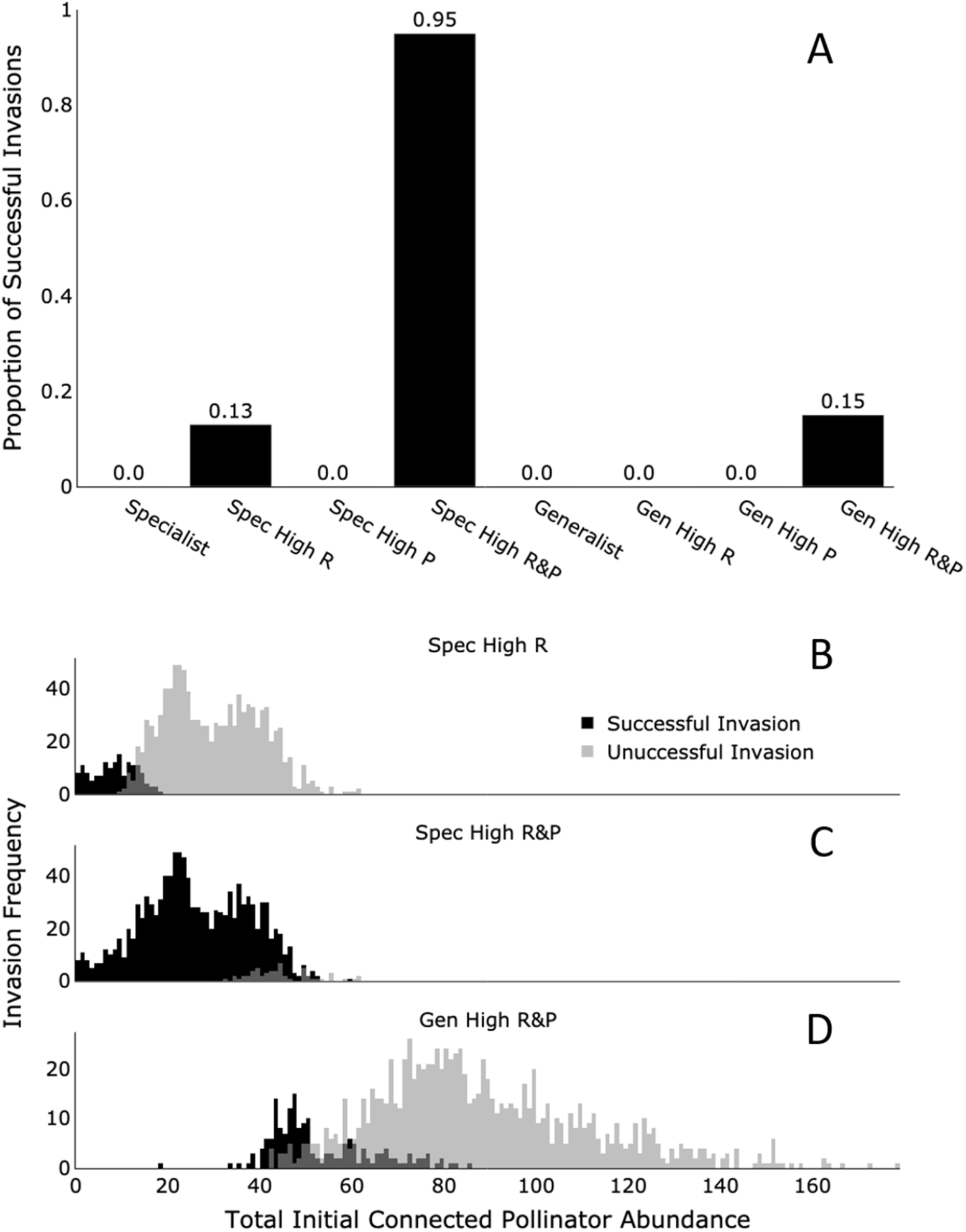
Proportion of successful plant invasions of each introduced species type (A) and the effect of pollinator abundance initially visiting them on their invasion success (B-D). Panel **A** shows (N = 28,800) that introduced plants visited by one or a few native pollinator species (Spec), high reward producers (High R), and with high pollen attachability (High P) most frequently invaded. Introduced plants visited by many different pollinator species (Gen) and exhibiting the average level of rewards production or pollen attachability found among native plants (indicated by omitting High R or P) never invade. Panels **B, C, D** show data (N = 3,600; per panel) for the only three species types that successfully invaded the networks, that is, specialist plant species with high production of rewards (Spec High R), specialist plant species with high production of rewards and pollen attachability (Spec High R&P), and generalist plant species with high production of rewards and pollen attachability (Gen High R&P), respectively. Black and light gray bars represent successful and unsuccessful invasion, respectively, while medium gray indicates where those two bar types overlap.

**Table 4.** Classification and Regression Tree (CART) analyses for invasion success.

|  | Initial analysis | Refined analysis* |
| --- | --- | --- |
| **Five fold R<sup>2</sup>** | 0.82 | 0.87 |
| **Main Contributions** | High reward producer (34%)<br>More specialized (25%)<br>High pollen attachability (22%)<br>Linkage algorithm (5%) | High reward producer (36%)<br>*Initial pollinator abundance connected to non-native (33%)<br>High pollen attachability (31%) |

### How does the quantity and quality of visits a plant receives from resident pollinators affect their invasion success?

We found that plants visited by fewer pollinators (in terms of abundance) at the moment of their introduction were most likely to invade (Fig. 1B-C). Therefore, we conducted a second (refined, see Table 4) CART analysis in which we incorporated the initial pollinator abundance connected to the introduced plant as a contributor for the analysis. This refined analysis shows that the total abundance of pollinators visiting the introduced plant species better predicts its invasion success than the number of pollinator species visiting it (note these two variables are strongly and positively correlated, see Fig. S1).

The explanation for introduced plants visited by fewer pollinators being more likely to invade resides in the reward threshold determining whether a plant species attracts sustained visitation or not (hereafter “reward threshold”; Fig 2, Appendix S1, Fig. S2). When the reward density of a plant species drops from such threshold, the pollinators stop visiting it and the plant species declines in abundance which, in turn, reduces the reward density of its population even further (i.e., fewer flowers available for pollinators). This vicious cycle causes the irreversible process of plant species going extinct once their rewards density drops below the reward threshold. All plant species have the same reward threshold at each simulation (Eq. S2 in Appendix S1, R* in Fig S2), as a result of the “ideal-free distribution” caused by pollinators being adaptive foragers (Valdovinos et al. 2013), and its value is determined by the parameter values drawn randomly prior to running each simulation. However, the dynamics of floral rewards differ among plant species given that they have different per-capita production rate of rewards and are visited by different pollinator species with different abundances and foraging efforts.

**Figure 2.**
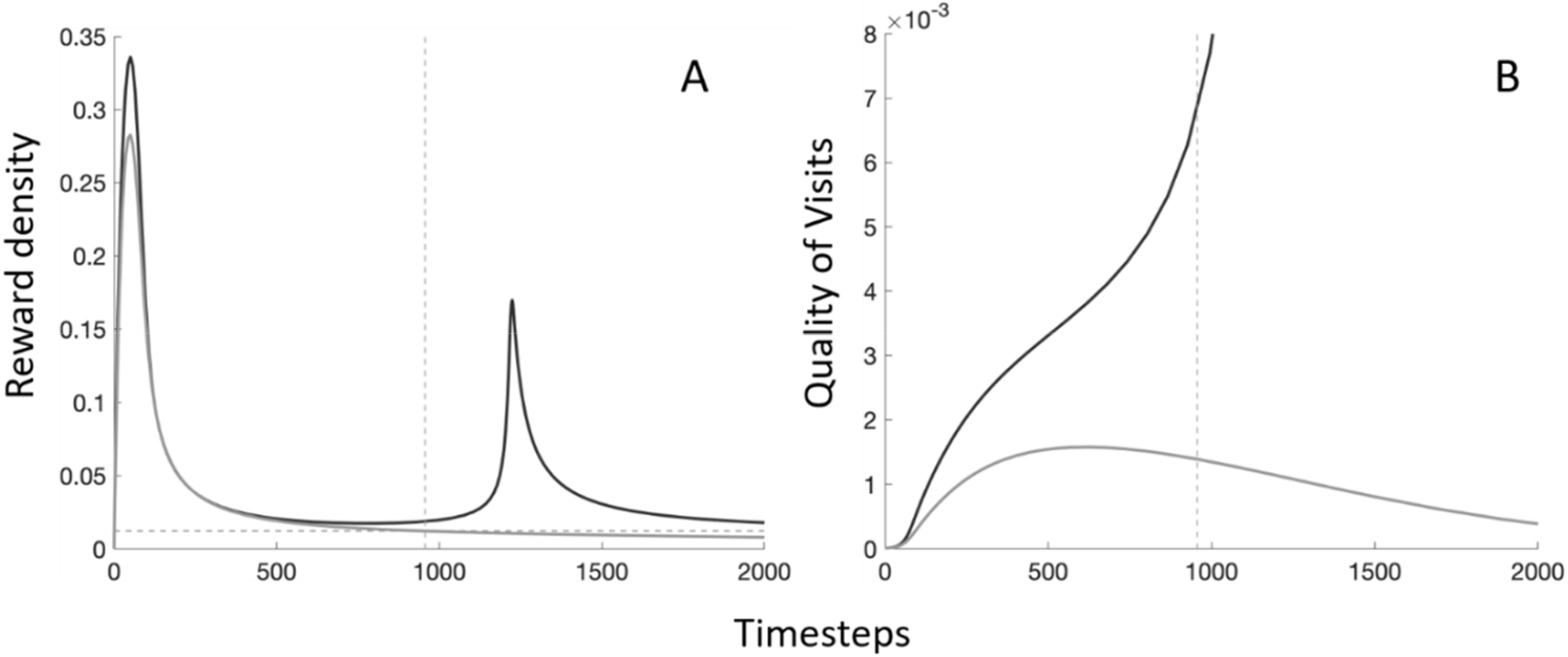
Reward threshold that determines invasion success during the transient dynamics. Transient dynamics are defined as the non-asymptotic dynamical regimes that persist for less than one to ‘as many as tens of generations’ (Hastings et al. 2018). Two simulations (one of the successful, black curves, and one of the failed, gray curves, invasions) for the introduction of specialist plant species with high production of rewards and pollen attachability (Spec High R&P) chosen from the data shown in Fig. 1C, to illustrate: **A**. An introduced plant species fails to invade (gray curve) when its rewards drop from the reward threshold (horizontal dashed line). The vertical dashed line indicates the timestep at which the reward threshold was crossed for the failed invasion. **B**. The quality of visits received by the introduced plant species does not increase enough for the failed invasion before the reward threshold is reached, so it goes extinct (see Fig S2). In the successful invasion, the introduced plant species is able to attract the pollinators’ foraging effort fast enough during the transient dynamics that it obtains enough quality of visits to persist before the threshold is met. The second peak observed in panel A corresponds to the increased floral rewards due to the increase in abundance of the introduced species that successfully invades, but then get depleted again to the reward density determining the system’s equilibrium (see Eq. S2 in Appendix S1). All successful and failed invasions look qualitatively the same as these figures.

If the reward density of the introduced species (black curve in Fig. 2A) stays at or above this reward threshold (grey dashed curve in Fig. 2A) the plant population keeps attracting pollinators for long enough to receive high quality of visits (black curve in Fig. 2B), which ensures its population growth and, therefore, its invasion success (Figs. S3A-D). If the reward density of the introduced species (grey curve in Fig. 2A) drops from this threshold due to high consumption by pollinators, the pollinators stop visiting it and reassign their foraging effort to other plant species in their diet whose rewards are at or above the threshold. Consequently, the plant species receives low-quality visits and goes extinct (compare gray with black curve in Fig 2B; Fig. S2). See Appendix S1 for a mathematical analysis demonstrating that our results on transient reward dynamics are general (hold true) across parameter values, which is stronger than conducting sensitivity analyses.

### How do plant invasions impact network structure and the reproduction success of native plants?

We found that the native plants that shared pollinator species with the successful invaders received lower quantity (Figs. 3A and 4A) and quality (Figs. 3B and 4B) of visits after the plant invasion, which is explained by pollinators re-assigning their foraging efforts from the native to the invasive plant species (Fig. 4D). However, the native plants only slightly decreased their density (Fig. 4C) and never went extinct (data not shown) as a consequence of the invasion. The magnitude of this negative effect on the density of native plants was reduced by the number of plant species in the network (Fig. 4G). Conversely, the plant invasions increased the density of native pollinators (Fig. 4F), an effect that was also attenuated by the number of plant species in the network (Fig. 4H). Finally, the plant invasions slightly increased the networks’ weighted nestedness (Fig. 3C) and modularity (Fig. 3D). See Table S1 for all the statistics of the Welch Two Sample t-test comparing weighted nestedness and modularity for all networks, groups of networks, and by the plant types introduced. Table S2 conceptually summarizes Table S1 for easy understanding of the trends.

**Figure 3.**
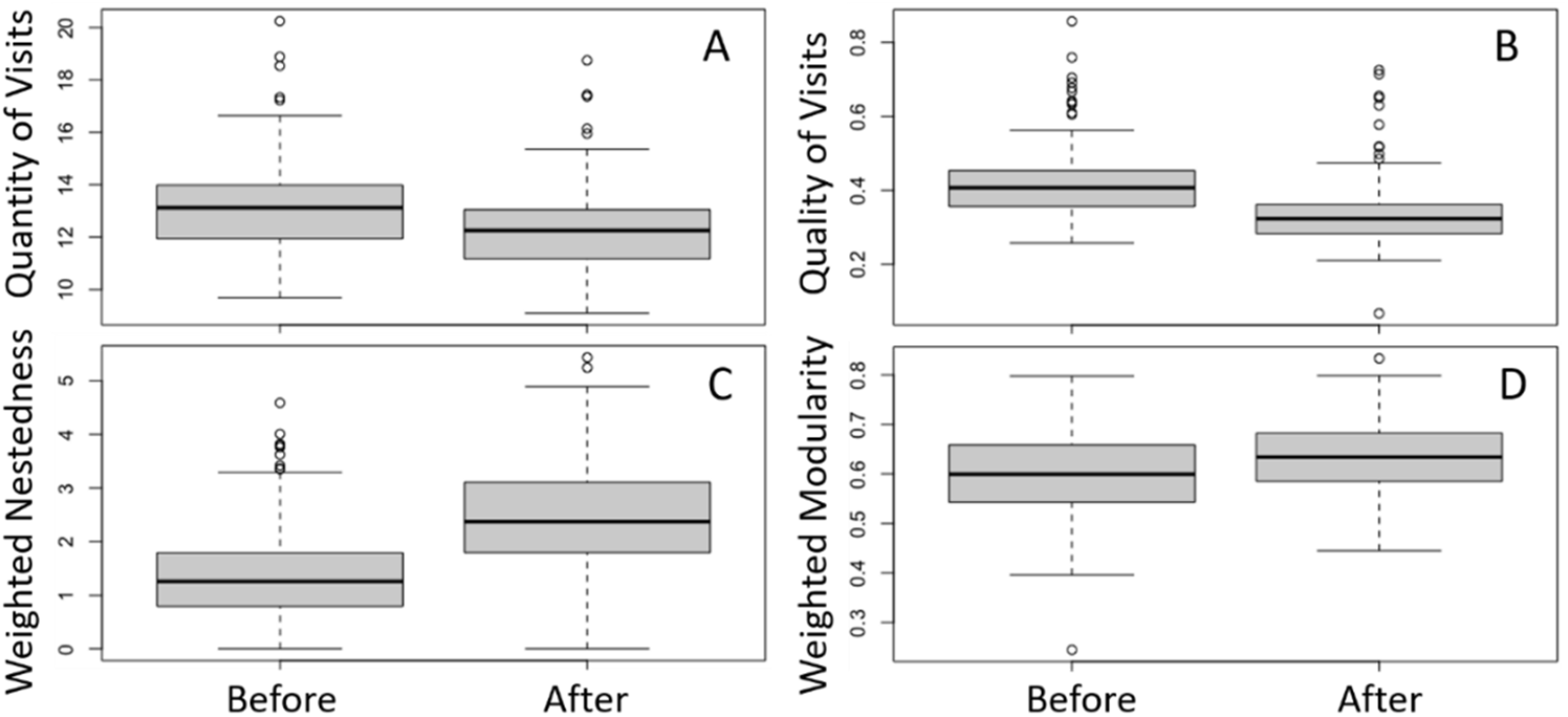
Effect of plant invasions on the quantity (A) and quality (B) of visits received by native plants and the networks’ weighted nestedness (C) and modularity (D). Box plots for these variables before (at 10,000 timesteps) and after (at 20,000 timesteps) the plant introduction for all the networks with 40 species and connectance 0.25 that were invaded by the three plant types that successfully invaded the networks (see Fig 1A). The middle bar, box, and error bars represent the mean, interquartile range, and standard deviations of each distribution. Welch Two Sample t-test for A, B, C, and D show significant differences between the variable means before and after invasion, all of which generated p-values less than 10^−7^ (see Table S2). We found a negative correlation between weighted nestedness and modularity (Fig. S5A; correlation coefficient -0.17 by Pearson’s test) – consistent with previous analysis on binary structure (Fortuna et al. 2010) – which became more negative after the invasion (Fig. S5B; correlation coefficient -0.50). See Fig. S6 showing the same qualitative results of panels C and D but when the invader and their interactions are removed from the analyses of network structure after the invasion. That is, keeping network size and species composition constant before and after the invasion did not change our results.

**Figure 4:**
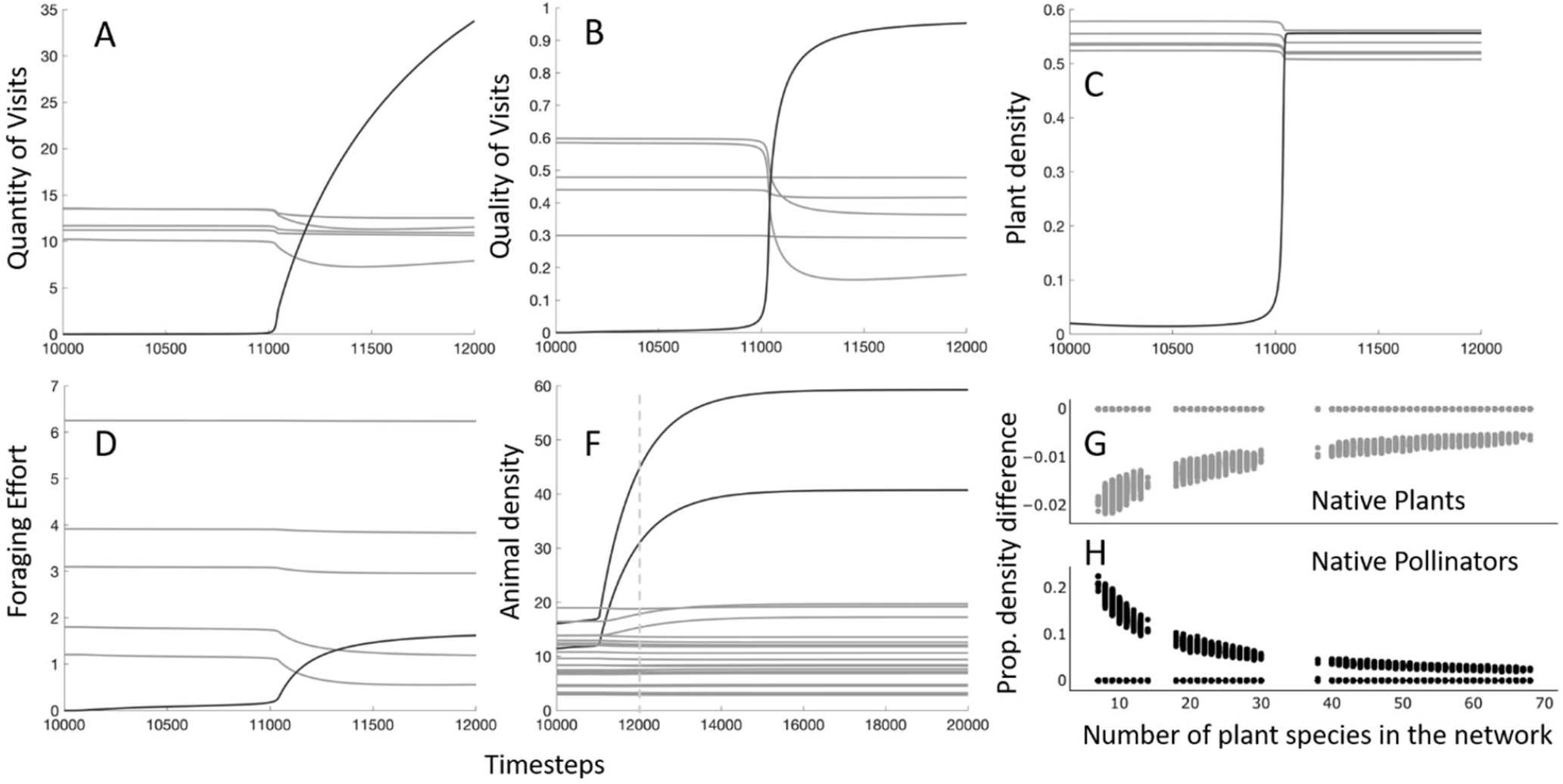
Effects of plant invasions on native plants (A-D, G) and pollinators (F and H) right after the plant introduction. Panels A-F show time series for only one simulation chosen from a successful invasion of Spec High R&P, but all simulations with successful invasions show qualitatively similar patterns. Quantity (A) and quality (B) of visits, density (C), and foraging effort assigned to the invasive plant species (black) increase over time, while those of native plant species (gray) sharing pollinators with the invasive species decrease. Panel F shows the increase in density of pollinator species (black) visiting the invasive species in comparison to those (gray) not visiting the invasive. Panels G-H show the results of all simulations in which specialist plant species with high production of rewards and pollen attachability (Spec High R&P) were introduced (Fig. 1C), with each dot representing one simulation. Plant richness decreases the magnitude of the negative (G) and positive (H) effects of the plant invasion on the native plants and pollinators, respectively, which is consistent with Elton’s (1958) prediction of richer systems being more robust to species invasions than poorer systems.

## Discussion

We found that 1) introduced plant species producing more floral rewards than natives were more likely to invade, 2) introduced species visited by fewer pollinators but receiving higher quality visits were more likely to invade, and 3) plant invasions decreased the quantity and quality of visits received by the native plants, slightly increased the network’s weighted nestedness and modularity, and slightly decreased the reproduction success of native plants.

Network structure did not predict the plant invasion success (results 1 and 2) but affected the impacts on natives (result 3); that is, the number of plant species in the network decreased the magnitude of the invaders’ negative and positive effects on native plants and pollinators, respectively.

Our first two results are a consequence of the transient dynamics that occur right after the plant introduction. These dynamics occur because plants are introduced at very low abundances (Appendix S1) so they need to produce more rewards than the natives to attract pollinators.

Introduced plants need those pollinators to increase their foraging effort by a great amount for them to become efficient (i.e., carrying mostly the conspecific pollen of the introduced plant). Receiving visits by many pollinator species or by abundant pollinators depletes the rewards of the introduced plant more quickly to the reward threshold that determines the system’s equilibrium. Therefore, pollinators stop reassigning their foraging effort to the introduced plant before they become efficient pollinators and the introduced plant goes extinct. To the best of our knowledge, our work is one of the first revealing a dynamical transient in ecological networks, as theory on ecological networks has traditionally focused on equilibrium dynamics (e.g., Bascompte et al. 2006, Bastolla et al. 2009, Pascual-García and Bastolla 2017, Valdovinos and Marsland 2020).

Mathematical discussion of the importance of transients traditionally takes place in the context of systems where the fixed point is never reached (whether due to limit cycles, chaos, stochastic perturbations, etc.), or where the time scale for equilibration is so long that the fixed point is irrelevant (Hastings et al. 2018, 2021). However, our results demonstrate that while there is always a stable fixed point in which non-native plant species invade, the ability for the system to reach that point from the initial conditions of low non-native plant abundance is based on the transient dynamics of reward density. Specifically, we found that based on the rate at which non-native plant species’ rewards are reduced to equilibrium, they either secure sufficiently efficient visits to invade or do not and go extinct. We show in Fig. S4 that increasing the initial abundance of non-native species 10 times, which increases their population reward production by 10 times, allows all plant types to invade including the generalists. This suggests that there is some reward production level that always produces a successful invasion (given a fixed native community) with a sharp threshold separating from the region of failed invasion. Future mathematical work should analyze this tipping point by finding the threshold in initial plant abundance, and therefore reward production, determining plant invasion success.

Our finding of higher invasion success of plants offering higher amounts of floral rewards is consistent with empirical research showing that plants that successfully invade plant-pollinator networks typically offer large amounts of floral rewards in large, showy flowers (Lopezaraiza–Mikel et al. 2007, Muñoz and Cavieres 2008, Padrón et al. 2009, Pyšek et al. 2011, Kaiser-Bunbury et al. 2011). Empirical data also support our findings that plant invasions can increase the abundance of native pollinators (Lopezaraiza–Mikel et al. 2007, Bartomeus et al. 2008, Carvalheiro et al. 2008), but decrease the quantity and quality of visits received by native plants (Traveset and Richardson 2006, 2014, Morales and Traveset 2009, Arceo-Gómez and Ashman 2016, Kaiser-Bunbury et al. 2017, Parra-Tabla et al. 2021). Finally, we found that plant invasions made the network structures slightly more nested and modular, which is consistent with previous theoretical (Valdovinos et al. 2009) and empirical (Bartomeus et al. 2008) work, respectively. Other empirical studies did not find a clear difference in structure between invaded and uninvaded networks (Vilà et al. 2009, Albrecht et al. 2014, Parra-Tabla et al. 2019). The field, however, still lacks understanding on how those effects of invasive plants on visitation rates and network structure translate to effects on the reproduction success and population growth of native plants (Parra-Tabla and Arceo-Gómez 2021). Our work can help guide future empirical research by showing that when other stages of plant reproduction are considered beyond visitation (i.e., successful pollination events, seed production, recruitment), a decrease in quantity or quality of visits does not necessarily translate into a decrease in plant reproduction or reduction of plant growth.

We found no extinction caused by the plant invaders, which is explained by: 1) plants only needing a few high-quality visits to produce enough seeds, and 2) seed recruitment being dependent on competition among plants for resources other than pollinators, with intraspecific stronger than interspecific competition (see Table 1). Native plants receive enough high-quality visits before the plant introduction and grow in abundance up to their equilibrium point determined mostly by their intraspecific competition (Valdovinos and Marsland 2020). The reduction of adaptive foraging reallocated from the native to the non-native plants is always smaller than what would be needed for the native plant to receive sufficiently low-quality visits to be driven extinct. Therefore, our work suggests that competition for pollinators alone is not enough to cause native plant extinctions. Future work should evaluate how competition between natives and invaders for resources other than pollinators affect the persistence of native plant species (Mitchell et al. 2006).

Our study is limited to the analysis of non-native plants that are completely dependent on pollinators to persist and that are introduced only once and in very small numbers. Regarding the first limitation, successfully invading plants are often not completely dependent on animal pollinators for reproduction, with many being abiotically pollinated or capable of some level of autogamous selfing or asexual reproduction (Barrett 2011, Burns et al. 2011). Second, introducing plants only once and in very small numbers is at the core of our results showing that generalist plants are less successful at invading networks than specialist plants. In fact, increasing their initial abundance 10 times – as mentioned above – allowed all generalist types to invade (Fig. S4A). Our results suggest that the common finding of invasive species often exhibiting “highly generalized floral traits” (e.g., radial symmetry; reviewed in Parra-Tabla and Arceo-Gómez 2021) might be explained by those taxa being introduced several times and at larger numbers than those we simulated here.

Finally, to our knowledge, ours is the first study suggesting that the cost of too many visits can affect the invasion success of non-native plants. This initial introduction process into plant-pollinator networks is difficult to study empirically because it would require conducting the study during the first arrival of the non-native plant, or deliberately introducing the plants, which poses ethical problems (Stricker et al. 2015). Therefore, our study also exemplifies how theoretical work can promote new thinking and research in areas traditionally studied empirically. Overall, our work contributes in promoting new thinking to integrate theoretical and empirical research during the transient dynamics of ecological networks, and calls for evaluating the impact of invasive plants not only on visitation rates and network structure, but also on the demographics of native plants, which depend on other processes beyond animal visitation such as seed production and recruitment.

## Supporting information

All supplementary information in one file

