## Supplementary material for "Transient dynamics in plant-pollinator networks: Fewer but higher quality of pollinator visits determines plant invasion success": All supplementary information in one file

Supplementary Information, in the order that it is first mentioned in the main text, includes:

**Appendix S1:** Mathematical analysis confirming the generality of our simulation results. We provide mathematical relationships between our model's parameters, variables, and invasion conditions that determine invasion success.

**Table S1:** Network structure properties and non-native traits totaled 21 contributors for our Classification and Regression Tree (CART) analysis.

**Figure S1:** Total abundance (density) of pollinators initially visiting the introduced plant species regressed against the number of pollinator species initially visiting the plant species, for the three linkage algorithms.

**Figure S2:** Impact vectors and zero net growth isoclines (ZNGIs) from Valdovinos and Marsland (2021) illustrating how much the ZNGIs need to move due to adaptive foraging for the quality of visits to improve.

**Figure S3:** Time-series of reward density, foraging effort, quality of visits, and density of an introduced plant species that successfully invaded the network and one that failed to invade.

**Figure S4:** Increasing introduced plant abundance from 0.02 (value used in the main text, which is the plant extinction threshold in the model) to 0.2 allowed all introduced plant species to successfully invade. The positive and negative impacts of the invaders on native pollinator and plant species, respectively, were equivalent to those exerted by invaders introduced at 0.02.

**Figure S5:** Negative correlation between weighted nestedness and weighted modularity before and after the plant invasion. These metrics were calculated for all the networks with 40 species and connectance 0.25 that were successfully invaded.

**Figure S6:** Effect of plant invasions on the networks' weighted nestedness (A) and modularity (B) when the invader and its interactions are excluded from the analysis, as opposed to Figures 3C and 3D of the main text, respectively, which include the invader and its interaction for calculating the network metrics after the invasion.

**Table S2:** Statistics for Welch Two Sample t-test comparing the distributions of weighted nesteness and weighted modularity of networks before and after the plant invasions.

**Table S3:** Conceptual summary of Table S2 for fast and easy understanding of our results on the effect of plant invasions on the networks' weighted structure.

**Appendix S1:** Mathematical analysis confirming the generality of our simulation results

Valdovinos and Marsland (2021) show mathematically that the key determinant of plant survival in Valdovinos et al.'s (2013) model is the quality of visits they receive, $\sigma_{ij} = \frac{\varepsilon_i \alpha_{ij} p_i}{\sum_{k \in P_j} \varepsilon_k \alpha_{kj} p_k}$ (see parameter and variable definitions in Table 1 of main text). Moreover, the authors find a threshold of quality of visits that plants need to meet to persist and the two equilibrium conditions determining such threshold (Eqs. S1 and S2). We use these two conditions to determine when an introduced plant species ($i = x$) will invade the network. First, the rewards at equilibrium of the introduced plant species $x$ is obtained by solving $\frac{dR_x}{dt} = 0$ (Eq. 3 in main text), which results in:

$$R_x = \frac{\beta_x p_x}{\varphi_x + \sum_{j \in A_x} \beta_x \tau_{xj} \alpha_{xj} a_j} \quad \text{(Eq. S1)}$$

This condition indicates that the floral rewards at equilibrium of the introduced plant species is higher when the plant has a higher per-capita production rate $\beta_x$, and decreases with the total abundance of pollinators visiting the introduced plant $\sum_{j \in A_x} \alpha_{xj} a_j$. The latter expression is key to our analysis, as it shows that highly abundant pollinators (i.e., high $a_j$) need to increase less their foraging effort to the introduced plant (i.e., low increase of $\alpha_{xj}$) to decrease its rewards to levels where the entire system (i.e., all species abundances, rewards, and foraging efforts) equilibrates (see Eq. S2 and Fig. S2).

Second, the abundance of pollinators visiting the introduced plant species $x$ and their foraging efforts also reach an equilibrium, which Valdovinos and Marsland (2021) showed is guaranteed by:

$$R_x = R_x^* = \frac{\mu_x^A}{c_{xj} b_{xj} \tau_j} \quad \text{(Eq. S2)}$$

This condition together with Eq. S1 indicate that the system reaches a new equilibrium that includes the new plant species when the consumption of the rewards of $x$ by the native pollinators drive the abundance of those rewards to $R_x^*$. If the rewards of $x$ drop below $R_x^*$, the native pollinators will re-assign their foraging efforts to the native plants and stop visiting the introduced species. Importantly, these conditions are general and make our mathematical analysis robust to all parameter values, network structures, and linkage algorithms.

Our mathematical analysis explains our simulation results and demonstrate that they are robust. Eq. S1 shows that the higher the production of rewards of the introduced plant ($\beta_x$), the higher the amount by which the pollinators' foraging efforts ($\alpha_{xj}$) assigned to the non-native will increase and, therefore, more likely is the non-native plant to invade because it will receive higher quality of visits ($\sigma_{ij}$). This explains why high rewards producers can invade.

Inspection of the key term $\sum_{j \in A_x} \alpha_{xj} a_j$ in Eq. S1 also explains the negative relation between abundance of pollinators initially visiting the non-native plant and its invasion success (Fig. 1B-D). This term indicates that the more abundant the pollinators visiting the introduced plant species initially ($\sum_{j \in A_x} a_j$) the less their foraging effort assigned to the introduced plant ($\alpha_{xj}$) need to increase to decrease the introduced plant's rewards to the system's equilibrium specified by Eq. S2 (where the rewards of the introduced species, the foraging efforts of pollinators, and the pollinator abundances stabilize). When this increase in foraging efforts is not enough for the introduced species to receive the quality of visits needed to invade (see variable $\alpha_{ij}$ in the numerator of $\sigma_{ij}$, above; Fig. S2), the introduced plant goes extinct.

Finally, note that invader reward $R_x$ must exceed equilibrated native rewards $R_i^*$ for long enough to attract sufficient pollinators, which value (using parameter values from Table 2) is:

$$R_i^* = \frac{\mu_j^A}{c_{ij} b_{ij} \tau_j} = \frac{1}{80} = 0.0125 \quad \text{(Eq. S3)}$$

**Table S1:** Network structure properties and non-native traits totaled 21 contributors for our Classification and Regression Tree (CART) analysis. Non-native plant properties included the generality level (Inv_spec), pollen attachability (Inv_pol), rewards production (Inv_rew), and the linkage algorithm (Inv_k_alg). Network structure properties included species richness (*S*), the ratio of animal to plant species (A/P), four measures of link density [connectance ($C = L$ / A×P, where *L* is the total number of links, A the number of pollinator species, and *P* the number of plant species), links per species (L/S), links per plant species (L/P), and links per animal species (*L/A*)], four measures of degree distribution [power law exponent for plants (*gP*) and animals (*gA*)], the standard deviation of animal generality (GenSD) and the standard deviation of plant vulnerability (VulSD)], four measures of niche overlap [the mean and maximum Jaccardian index for plants (mJP, maxJP) and animals (mJA, maxJA)], and nestedness (NODFst). Further description of these properties can be found in Valdovinos et al (2018, *Nature Communications*) and reference therein.

**Column Contributions**

| Term | Number of Splits | G^2 | Portion |
|---|---|---|---|
| InvP_rew | 1 | 6947.62765 | 0.3417 |
| InvP_spec | 3 | 5181.28023 | 0.2548 |
| InvP_pol | 1 | 4506.63184 | 0.2216 |
| InvP_k_alg | 11 | 1049.4083 | 0.0516 |
| NODFst | 7 | 910.624617 | 0.0448 |
| C | 3 | 788.41759 | 0.0388 |
| VulSD | 2 | 176.148891 | 0.0087 |
| maxJP | 3 | 174.217863 | 0.0086 |
| S | 1 | 145.455931 | 0.0072 |
| L/A | 3 | 101.838225 | 0.0050 |
| mJA | 4 | 96.8494324 | 0.0048 |
| L/P | 3 | 76.5449778 | 0.0038 |
| L/S | 3 | 74.7496029 | 0.0037 |
| A/P | 2 | 42.631865 | 0.0021 |
| GenSD | 2 | 34.1652958 | 0.0017 |
| mJP | 1 | 13.5058517 | 0.0007 |
| gP | 1 | 13.2094611 | 0.0006 |
| P | 0 | 0 | 0.0000 |
| A | 0 | 0 | 0.0000 |
| gA | 0 | 0 | 0.0000 |
| maxJA | 0 | 0 | 0.0000 |

**Fig. S1.**

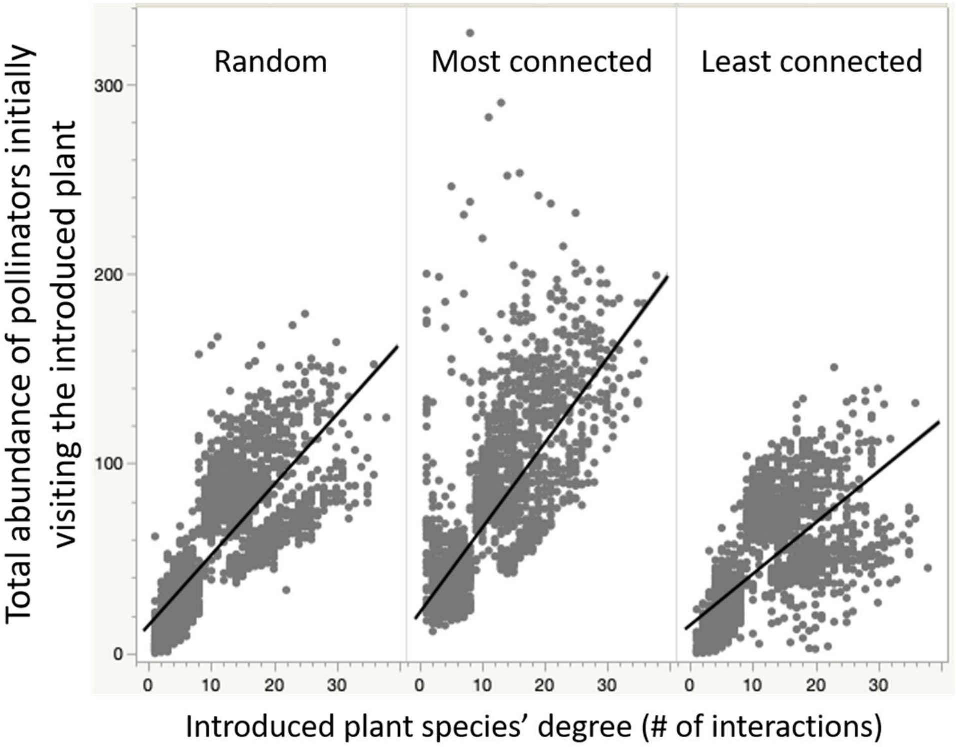


**Figure S1. Regressions showing the strong positive relation between the number of pollinator species initially visiting the introduced plant species (x-axes) and the total abundance of those pollinators (y-axis).** The three panels show data for the three linkage algorithms (see Methods in main text): (1) random, in which the introduced plant species was connected randomly to any pollinator species in the network, (2) most connected, in which the introduced plant species was connected randomly to the most-generalist pollinator species (i.e., those visiting the highest number of plant species in the network), (3) least connected, in which the introduced plant species was connected randomly to the most-specialist pollinator species (i.e., those visiting the lowest number of plant species in the network). The regression statistics for each of these panels are 0.761 (for random), 0.712 (for most connected), and 0.659 (for least connected), with all p-values lower than $10^{-7}$. The regression statistics for all data combined are 0.674 with p-value lower than $10^{-7}$.

**Fig. S2.**

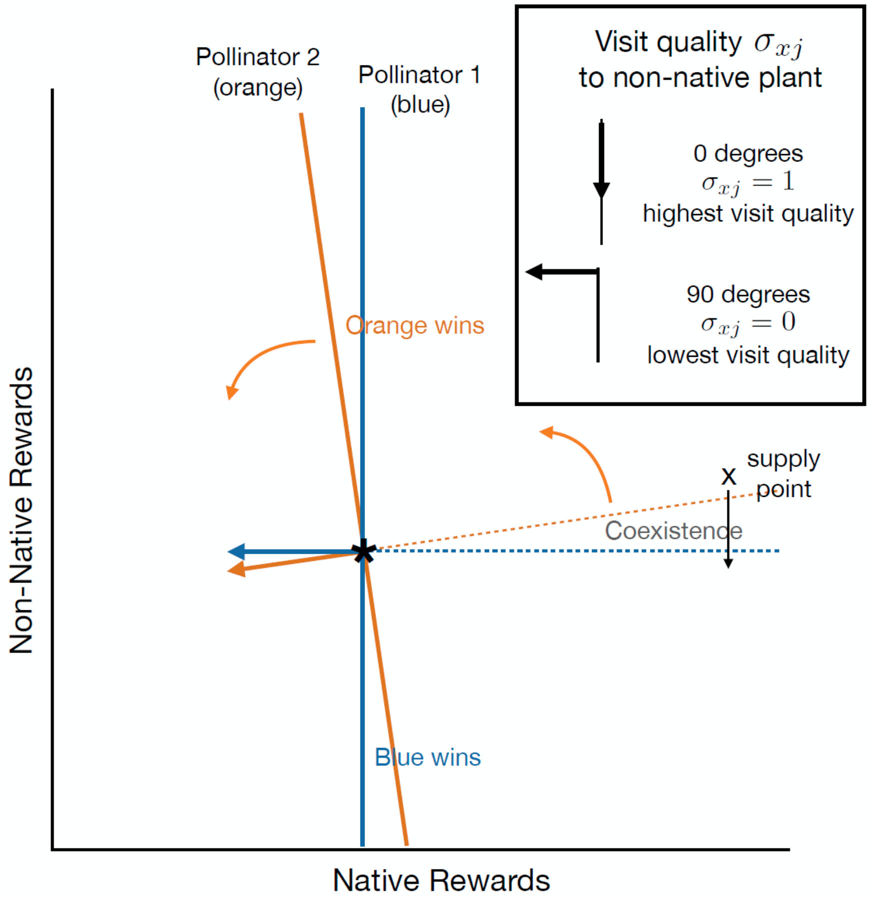


**Figure S2.** Impact vectors and zero net growth isoclines (ZNGIs) are shown for two pollinator species (blue and orange) competing for the rewards of two plant species, as in Valdovinos and Marsland (2021). Here, only the orange pollinator species is capable of foraging on the newly introduced plant species. Adaptive foraging causes the orange ZNGI and impact vectors to rotate in the direction of the most abundant rewards. The angle between the impact vector and the non-native rewards axis affects the visit quality, which improves as the angle decreases from 90 degrees ($\sigma_{ij} = 0$, lowest visit quality) to 0 degrees ($\sigma_{ij} = 1$, highest visit quality). When the non-native rewards level is depleted to the ideal-free distribution level R* (black star), the plants become equivalent in value for the pollinator, and adaptation stops. A higher visit quality can be achieved as adaptation continues during the transient characterized by the animal population requiring a large time interval in order to grow to the population size required to balance the supply. In principle, this rewards level can be stably achieved as soon as the supply point enters the coexistence cone, through the combined effects of adaptive foraging that expands the cone and decreases non-native rewards (i.e., moves the supply point down). But if the pollinator population depletes the non-native rewards below R*, they will stop visiting the non-native species which will result in the non-native extinction.

**Fig. S3.**

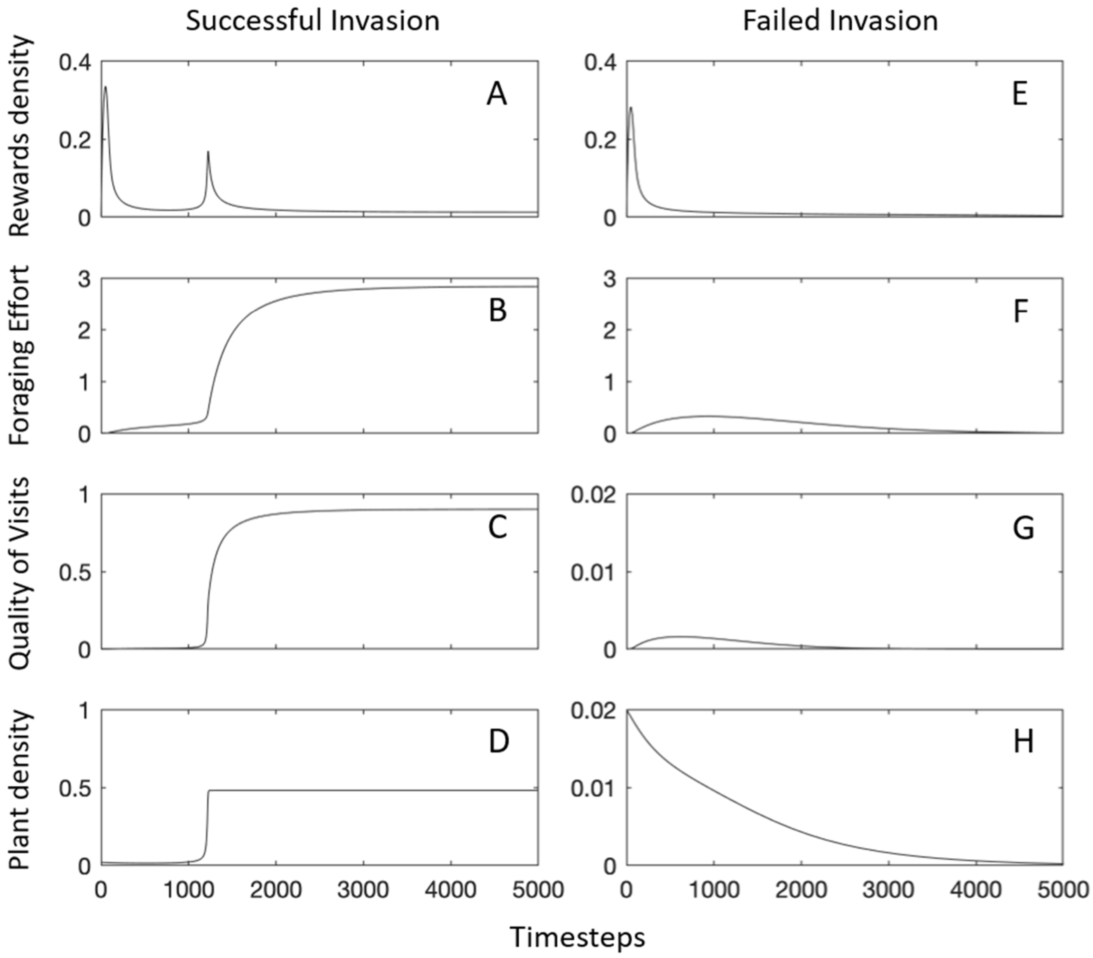


**Figure S3. Time-series of key variables indicating a successful (A-D) or failed (E-H) invasion.** Two simulations chosen from the data shown in Fig. 1C of the main text by the same type of introduced plant (specialist high rewards and pollen producer) and same network structure (40 species and connectance 0.25). This figure illustrates how an introduced plant species fails to invade when its rewards drop from the level at which the system reaches its feasible equilibrium (see Eq. 7 of main text), and complements with more variables the same case shown in Fig. 2 of the main text, which only shows reward density and quality of visits during the transient dynamics (i.e., only until 2,000 timesteps). All our successful and failed invasions look qualitatively the same as these figures, which we only use to illustrate our mathematical results that are general for all parameter values and network structures.

**Fig. S4.**

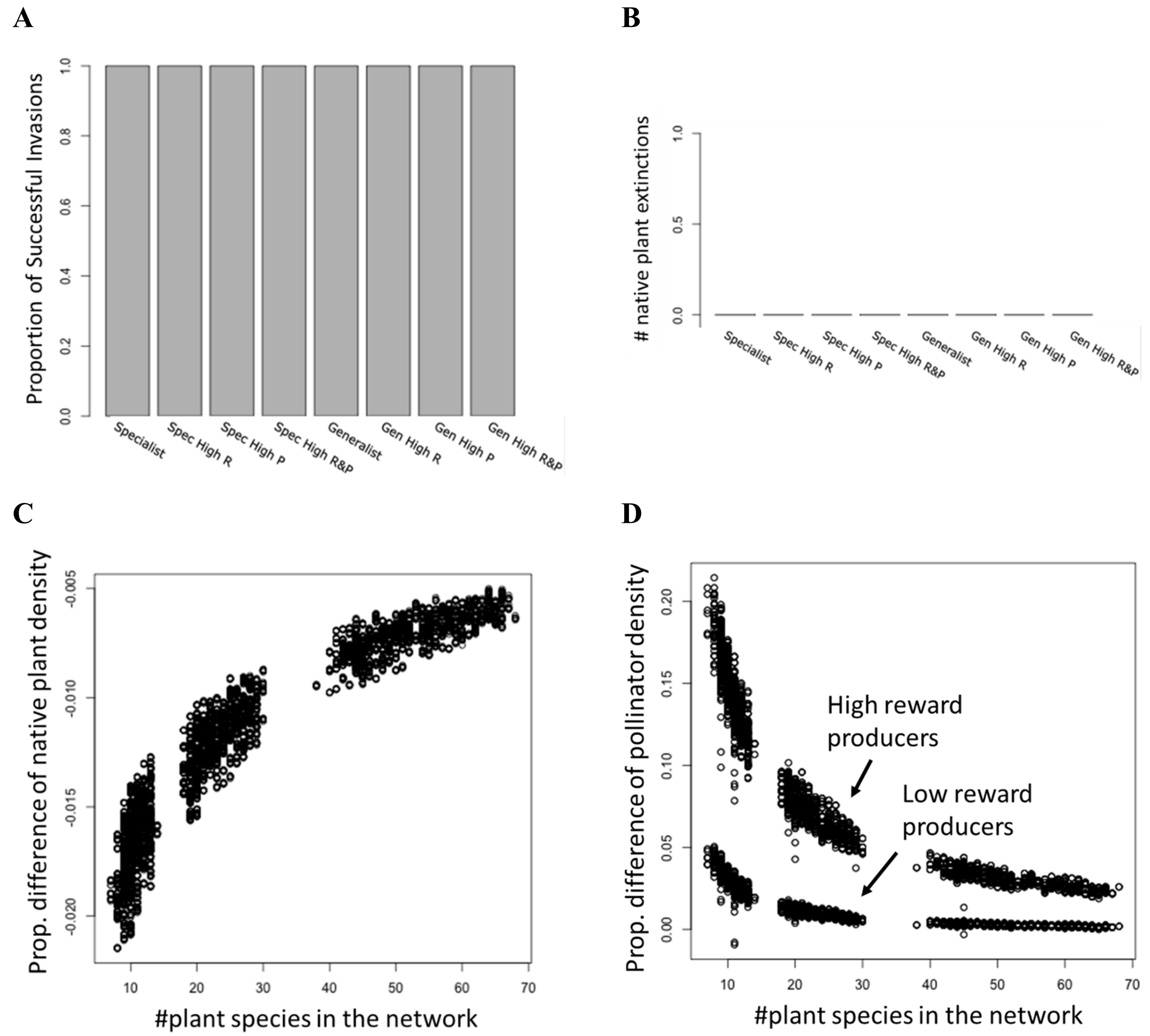


**Figure S4.** Results for increasing introduced plant abundance from 0.02 – value used in the main text that is also the plant extinction threshold in the model – to 0.2 individuals/area, which is still 4 times lower than the abundance at equilibrium of native plants (around 0.8 individuals/area). **(A)** All non-native plants invaded when introduced at such higher abundance. These invasions **(B)** caused no extinctions but **(C)** decreased the density of native plants and **(D)** increase the density of pollinators; both effects decreasing in magnitude with the number of plant species in the networks**.** Panels **A**, **C** and **D** show similar simulated introductions (N = 28,800) as the ones in Figures 1A, 4G, 4H of the main text. The greater positive effect on pollinators shown in Panel **D** is caused by invader types with high reward production (Spec High R, Spec High R&P, Gen High R, Gen High R&P), while the lower effects are caused by average reward production (Spec, Spec High P, Gen, Gen High P). Simulations in the main text (Figures 4H) do not have 2 sets of effects because average reward production plants do not invade. The negative and positive impacts of the high reward producer invaders on native plant and pollinator species, respectively, were equivalent to those exerted by the same invader types introduced at 0.02 (Figures 4G and 4H).

**Fig. S5.**

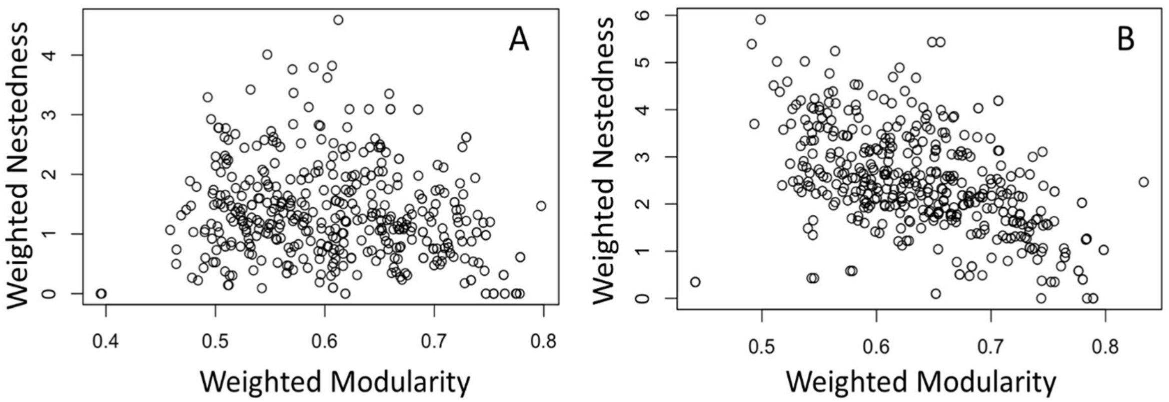


**Figure S5.** Correlation between weighted nestedness and weighted modularity before (**A**, at 10,000 timesteps; correlation coefficient -0.17) and after (**B**, at 20,000 timesteps; correlation coefficient -0.50) the invasion. These metrics were calculated for all the networks with 40 species and connectance 0.25 that were invaded by the three plant types that successfully invaded the networks (see Fig 1A). Correlation coefficients were calculated using Pearson's correlation test.

**Fig. S6.**

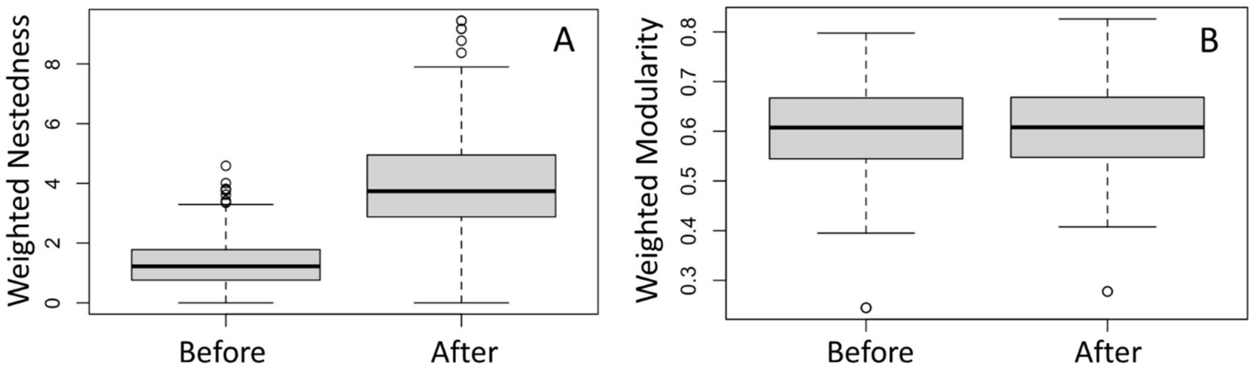


**Figure S6.** Effect of plant invasions on the networks' weighted nestedness (A) and modularity (B) when the invader and its interactions are excluded from the analysis, as opposed to Figures 3C and 3D of the main text, respectively, which include the invader and its interaction for calculating the network metrics after the invasion. Box plots for these variables before (at 10,000 timesteps) and after (at 20,000 timesteps) the plant introduction for all the networks with 40 species and connectance 0.25 that were invaded by the three plant types that successfully invaded the networks (see Fig 1A). The middle bar, box, and error bars represent the mean, interquartile range, and standard deviations of each distribution.

**Table S2:** Statistics for Welch Two Sample t-test comparing the distributions of quantity and quality of visits received by native plants and the networks' structure before and after successful plant invasions. The quantity and quality of visits were calculated only for native plant species that share a pollinator with the invasive plant species. Nestedness and modularity were measured on the quantitative network of visitation frequencies using the nest.smdm() and computeModules() functions, respectively, from the R package bipartite. Statistics were measured separately for the networks grouped by richness (S) and connectance (C) since those binary network properties influence the total number of visits in each network. Statistics were also measured separately for each invader type and for all invader types combined to determine whether the trends differed between plant types introduced. Differences in variable means before and after species invasions were analyzed with the Mann-Whitney test using the wilcox.test() function from the R package stats. This test was chosen because the assumption for equal variances required of a *t*-test was not always satisfied. Results from the Mann-Whitney test are less robust with smaller group sizes which was the case for results obtained by introducing "specialist plants with high production of rewards" (Spec High R, see main text) and "generalist plants with high production of rewards and high pollen attachability" (Gen High R&P).

| | | **First Set 400 Networks** *S* = 40 *C* = 0.25 | | | | **Second Set 400 Networks** *S* = 90 *C* = 0.15 | | | | **Third Set 400 Networks** *S* = 200 *C* = 0.06 | | | |
|---|---|---|---|---|---|---|---|---|---|---|---|---|---|
| **Invader Type** | **Variable** | **Initial Mean (SD)** | **Final Mean (SD)** | ***W*-statistic** | ***P*-value** | **Initial Mean (SD)** | **Final Mean (SD)** | ***W*-statistic** | ***P*-value** | **Initial Mean (SD)** | **Final Mean (SD)** | ***W*-statistic** | ***P*-value** |
| Spec High R | | n = 34 | | | | n = 30 | | | | n = 91 | | | |
| | Quality | 0.475 (0.134) | 0.409 (0.133) | 789 | 0.009 | 0.413 (0.136) | 0.362 (0.134) | 592 | 0.036 | 0.277 (0.081) | 0.245 (0.075) | 5203 | 0.003 |
| | Quantity | 13.704 (2.413) | 12.809 (2.210) | 730 | 0.063 | 6.496 (1.165) | 6.197 (1.101) | 521 | 0.300 | 2.194 (0.481) | 2.130 (0.461) | 4474 | 0.349 |
| | Nestedness | 0.695 (0.660) | 1.110 (0.871) | 404 | 0.033 | 0.321 (0.083) | 0.275 (0.072) | 405 | 0.513 | 1.256 (0.503) | 1.304 (0.501) | 3875 | 0.456 |
| | Modularity | 0.706 (0.096) | 0.730 (0.049) | 476 | 0.215 | 6.461 (0.991) | 6.156 (0.926) | 369 | 0.236 | 0.724 (0.062) | 0.735 (0.060) | 3606 | 0.133 |
| | | n = 362 | | | | n = 381 | | | | n = 400 | | | |
| Spec High P&R | Quality | 0.411 (0.078) | 0.330 (0.071) | 107073 | 2.2e-16 | 0.312 (0.073) | 0.268 (0.062) | 103435 | 2.2e-16 | 0.278 (0.056) | 0.249 (0.052) | 105962 | 2.0e-15 |
| | Quantity | 13.032 (1.518) | 12.178 (1.385) | 87651 | 3.7e-15 | 6.431 (0.972) | 6.129 (0.907) | 85216 | 3.2e-05 | 2.280 (0.468) | 2.214 (0.449) | 86391 | 0.051 |
| | Nestedness | 0.753 (0.508) | 1.495 (0.632) | 22022 | 2.2e-16 | 0.811 (0.473) | 1.104 (0.448) | 43215 | 2.2e-16 | 0.705 (0.535) | 0.771 (0.518) | 69583 | 0.001 |
| | Modularity | 0.597 (0.082) | 0.621 (0.066) | 52188 | 2.2e-06 | 0.571 (0.095) | 0.602 (0.085) | 55579 | 2.2e-08 | 0.604 (0.103) | 0.620 (0.097) | 70980 | 0.006 |
| Gen High P&R | | n = 16 | | | | n = 22 | | | | n = 141 | | | |
| | Quality | 0.453 (0.122) | 0.283 (0.033) | 254 | 1.3e-08 | 0.358 (0.076) | 0.276 (0.040) | 445 | 1.6e-07 | 0.306 (0.046) | 0.265 (0.037) | 15158 | 2.6e-14 |
| | Quantity | 13.402 (1.481) | 12.526 (1.356) | 174 | 0.086 | 6.923 (0.992) | 6.578 (0.933) | 296 | 0.212 | 2.383 (0.474) | 2.313 (0.455) | 10868 | 0.176 |

| | | | | | | | | | | | | | |
|---|---|---|---|---|---|---|---|---|---|---|---|---|---|
| | Nestedness | 0.504 (0.418) | 3.063 (1.131) | 15.5 | 2.4e-05 | 0.464 (0.392) | 1.304 (0.454) | 23 | 5.5e-09 | 0.392 (0.298) | 0.652 (0.409) | 3811 | 2.2e-16 |
| | Modularity | 0.558 (0.114) | 0.513 (0.055) | 190 | 0.019 | 0.564 (0.119) | 0.554 (0.100) | 243 | 0.991 | 0.557 (0.094) | 0.551 (0.079) | 9954 | 0.985 |
| All | | n = 412 | | | | n = 433 | | | | n = 632 | | | |
| | Quality | 0.417 (0.088) | 0.334 (0.080) | 137798 | 2.2e-16 | 0.321 (0.083) | 0.275 (0.072) | 133685 | 2.2e-16 | 0.284 (0.059) | 0.252 (0.053) | 267840 | 2.2e-16 |
| | Quantity | 13.102 (1.615) | 12.244 (1.476) | 112822 | 2.8e-16 | 6.461 (0.991) | 6.156 (0.926) | 109818 | 1.3e-05 | 2.291 (0.474) | 2.224 (0.455) | 215648 | 0.014 |
| | Nestedness | 0.739 (0.520) | 1.524 (0.752) | 30577 | 2.2e-16 | 0.808 (0.490) | 1.118 (0.469) | 55585 | 2.2e-16 | 0.714 (0.550) | 0.821 (0.533) | 161626 | 4.4e-09 |
| | Modularity | 0.605 (0.090) | 0.626 (0.074) | 71491 | 9.0e-05 | 0.583 (0.103) | 0.610 (0.094) | 75218 | 4.8e-07 | 0.611 (0.109) | 0.621 (0.104) | 185266 | 0.026 |

**Table S3:** Conceptual summary of Table S2 for fast and easy understanding of our results on the effect of plant invasions on the networks' weighted structure for each set of networks determined by their species richness (*S*) and connectance (*C*). Where * = <0.05, **=<0.001, ***=<0.0001.

| | Effect on Network | **First Set 400 Networks** $S$ = 40 $C$ = 0.25 | **Second Set 400 Networks** $S$ = 90 $C$ = 0.15 | **Third Set 400 Networks** $S$ = 200 $C$ = 0.06 |
|---|---|---|---|---|
| Specialist high reward producer | ↓ Quality | ** | * | ** |
| | ↓ Quantity | | | |
| | ↑ Nestedness | * | | |
| | ↑ Modularity | | | |
| Specialist high reward and pollen producer | ↓ Quality | *** | *** | *** |
| | ↓ Quantity | *** | *** | |
| | ↑ Nestedness | *** | *** | ** |
| | ↑ Modularity | *** | *** | * |
| Generalist high reward and pollen producer | ↓ Quality | *** | *** | *** |
| | ↓ Quantity | | | |
| | ↑ Nestedness | *** | *** | *** |
| | ↑ Modularity | | | |
| All Invader Types | ↓ Quality | *** | *** | *** |
| | ↓ Quantity | *** | *** | * |
| | ↑ Nestedness | *** | *** | *** |
| | ↑ Modularity | *** | *** | * |
